# Reverse genetics and comparative pathogenesis of Lone star virus

**DOI:** 10.64898/2026.07.06.736858

**Authors:** Dorcus C. A. Omoga, Curtis Witt, Henry Giesel, James M. Bowen, Krista Gunter, Stephanie Pozuelos, Ryan F. Relich, Benjamin Brennan, Natasha L. Tilston

**Author notes:** Corresponding author: Natasha L. Tilston (Tilston-Lunel).

## Abstract

Lone star virus (LSV) is a bandavirus first isolated from *Amblyomma americanum* ticks in the United States (U.S.) and is phylogenetically related to severe fever with thrombocytopenia syndrome virus (SFTSV), Heartland virus (HRTV), and Bhanja virus, each of which has been associated with severe human disease. In contrast to these better-characterized bandaviruses, LSV remains poorly studied, and its pathogenic potential is not well defined. Recent detection of LSV RNA in cerebrospinal fluid from an immunocompromised patient in Idaho, U.S., with fatal meningoencephalitis further highlights the need for experimental systems to investigate LSV biology. Here, we rescued recombinant (r) LSV from cloned cDNA and used it to characterize LSV. rLSV replicated similarly to the parental isolate in mammalian cells and caused rapid, systemic, and lethal disease in IFNAR^-/-^ mice, with widespread detection of viral (v) RNA across multiple tissues, hepatic and splenic pathology, and induction of inflammatory cytokines. In contrast, C57BL/6J mice controlled infection and exhibited no clinical disease. To place LSV within a broader comparative framework, we generated rSFTSV from cloned cDNA and compared rLSV, rSFTSV, and HRTV in cell culture and IFNAR^-/-^ mice. Our studies revealed distinct disease kinetics among these related tick-borne bandaviruses and showed that HRTV-induced immunity protected against homologous HRTV rechallenge and heterologous rSFTSV challenge, but not rLSV challenge. Together, these findings establish reverse-genetics platforms and small-animal models for comparative bandavirus studies, define key features of LSV pathogenesis, and place this neglected virus within a framework of related bandaviruses that differ in virulence and immunological overlap.

**Importance:** Tick-borne bandaviruses include several viruses associated with severe human disease, yet many related viruses remain poorly characterized. Lone star virus (LSV) was first isolated from *Amblyomma americanum* ticks decades ago, but experimental tools and animal models to study LSV infection have been lacking. Here, we generated a recombinant LSV and used it to define the outcome of infection in immunocompromised and immunocompetent mice. We show that LSV can cause rapid, systemic, and lethal disease when type I interferon signaling is absent, whereas immunocompetent mice restrict infection and remain clinically normal. By comparing LSV with Heartland virus and severe fever with thrombocytopenia syndrome virus, we also show that related tick-borne bandaviruses differ in disease kinetics and immune protection. These findings provide foundational tools for studying LSV and highlight the importance of experimentally characterizing neglected tick-borne viruses before their pathogenic potential is fully understood.

## Introduction

Tick-borne viruses represent a growing public health concern as changes in climate, land use, and human behavior drive the expansion of tick populations and increase human exposure^1,2^. Among these, tick-borne bunyaviruses, particularly members of the genus *Bandavirus* within the family *Phenuiviridae*, have emerged as important causes of severe human disease. Severe fever with thrombocytopenia syndrome virus (SFTSV) is now recognized as a major cause of hemorrhagic and neurological disease in East Asia, with case fatality rates reaching up to 30% in some outbreaks^3–10^. In the United States, Heartland virus (HRTV) has emerged as a tick-borne pathogen associated with severe febrile illness, cytopenias, neurologic complications, and fatal outcomes^11–18^. Bhanja virus (BHAV), another tick-borne bandavirus, has also been associated with febrile and neurologic disease in humans^19–22^. Together, these viruses underscore the pathogenic potential of tick-borne bandaviruses and highlight the need to better understand related but poorly characterized members of this group.

Lone star virus (LSV) was first isolated in 1967 from the Lone star tick, *Amblyomma americanum*, in the United States (U.S.)^23^, the same tick species implicated in HRTV transmission. Despite its early discovery and phylogenetic relatedness to HRTV, SFTSV, and BHAV, LSV has remained largely uncharacterized, and its pathogenic potential is unclear. Recently, LSV RNA was detected by metagenomic sequencing in the cerebrospinal fluid of a 60-year-old immunocompromised patient from Idaho, U.S., who developed progressive neurologic disease and fatal meningoencephalitis^24^. Although this single case does not establish causality or define the full clinical spectrum of LSV infection, it raises important questions about LSV virulence, tissue tropism, neurotropic potential, and host restriction.

Bandaviruses possess a tripartite RNA genome comprising a small (S) ambisense segment and medium (M) and large (L) negative-sense segments. The S segment encodes the nucleoprotein (N) and the nonstructural protein NSs in opposing orientations. The M segment encodes the glycoproteins Gn and Gc, and the L segment encodes the RNA-dependent RNA polymerase (RdRp). Each segment is flanked by untranslated regions (UTRs) that contain essential signals for viral transcription, genome replication, and packaging^9^. Studies of SFTSV and HRTV have demonstrated that type I interferon (IFN) responses play a central role in restricting bandavirus infection. In SFTSV, NSs potently suppresses innate immune signaling by sequestering key antiviral factors, including TBK1, IKKε, and IRF3^25–27^, whereas HRTV NSs employs a related but distinct strategy to interfere with TBK1–IRF3 signaling^14,28^. These virus-specific mechanisms of innate immune antagonism likely contribute to differences in replication, host restriction, and disease outcome among bandaviruses. However, whether LSV is similarly restricted by type I IFN responses and whether it causes systemic disease under conditions of impaired antiviral immunity has not been explored.

Reverse genetics systems have been essential for defining bandavirus biology and supporting countermeasure development^29–31^. For HRTV and SFTSV, recombinant virus systems have enabled direct interrogation of viral determinants of pathogenesis, including the role of NSs in innate immune antagonism and attenuation^7,8,28,32,33^. In contrast, comparable experimental tools and animal models for LSV are lacking, limiting similar studies. Under the institutional biosafety conditions used here in the U.S., LSV can be studied at biosafety level-2 (BSL-2), whereas HRTV and SFTSV require BSL-3 containment. Thus, a genetically tractable LSV system provides a complementary lower-containment platform for studying conserved and divergent features of tick-borne bandavirus biology.

In this study, we developed a reverse genetics system for LSV and used it to characterize LSV pathogenesis in mice. Using IFN alpha/beta receptor-knockout mice (IFNAR^⁻/⁻^) and immunocompetent wild-type (WT; C57BL/6J) mice, we demonstrate that LSV causes rapid, systemic, and lethal disease in the absence of type I IFN signaling, whereas infection is effectively restricted in immunocompetent hosts. We further compare LSV infection outcomes with those of HRTV and SFTSV, revealing marked differences in disease kinetics and immune protection among related tick-borne bandaviruses. Together, these findings establish LSV as a genetically tractable, BSL-2-accessible model for comparative bandavirus biology, define key features of LSV pathogenesis, and place this neglected virus within a framework of emerging tick-borne bandaviruses.

## Results

### Generation and characterization of a recombinant LSV

To establish a genetically tractable system for LSV, we generated a T7 polymerase-driven reverse genetics system using the LSV strain TMA1381. The parental virus, referred to here as WT LSV, was obtained from the World Reference Center for Emerging Viruses and Arboviruses [WRCEVA, University of Texas Medical Branch (UTMB)]. Published genome sequences for LSV strain TMA1381 are available in GenBank under accession numbers NC_021244, NC_021243, and NC_021242 for the S, M, and L segments, respectively. Sequencing of our WT LSV identified several nucleotide differences relative to the published TMA1381 reference sequence, including changes in the NSs, GPC, and RdRp coding regions (Fig. 1A).

**Figure 1.**
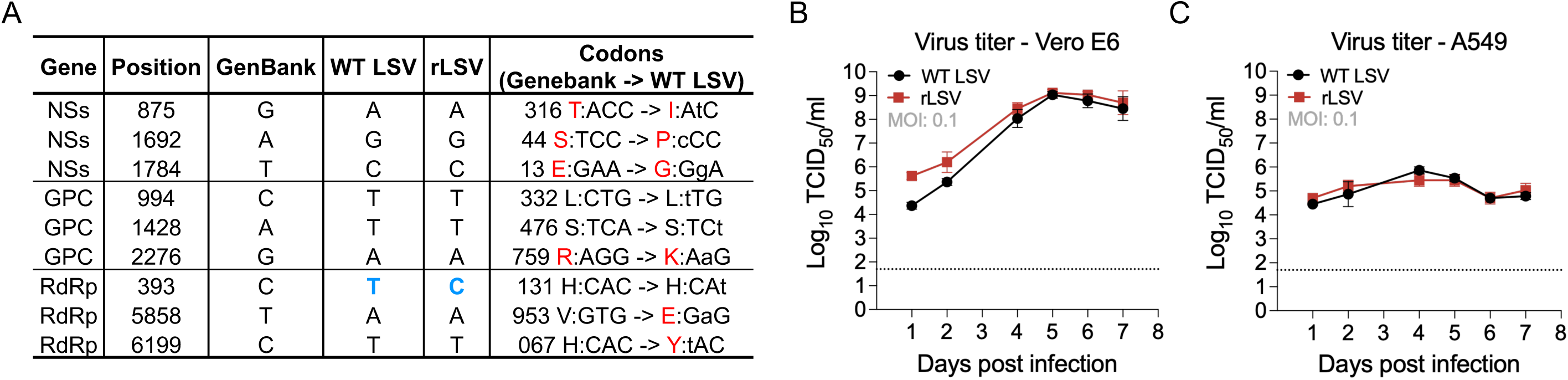
Characterization of rLSV. (A) Sequence comparison of WT LSV and rLSV relative to the published reference sequence (Accession number S: NC_021244, M: NC_021243, L: NC_021242). Nucleotide differences identified in the NSs, GPC, and RdRp coding regions are shown, together with the corresponding codon and amino acid changes. rLSV contained a single synonymous substitution relative to WT LSV used, highlighted in blue. (B, C) Multistep growth kinetics of WT LSV and rLSV in Vero E6 (B) and A549 (C) cells infected at an MOI of 0.1. Supernatants were collected at the indicated time points, and infectious titers were measured by a TCID_50_ assay. Data are shown as mean ± SD (n = 3). The dashed line indicates the limit of detection

Full-length cDNA copies of the S, M, and L segments were then generated from the sequenced WT LSV stock and cloned into T7-driven rescue plasmids. Co-transfection of the three rescue plasmids into BSR-T7/5 cells, together with helper plasmids expressing LSV N and RdRp, yielded infectious recombinant LSV (rLSV). The rescued virus was amplified in Vero E6 cells to generate working stocks reaching titers of up to 10^6^ TCID_50_/ml. Full-genome sequencing of rLSV identified a single synonymous substitution relative to the WT LSV stock used to generate the rescue plasmids, located at nucleotide position 393 in the RdRp ORF (T→C; His codon CAT→CAC; Fig. 1A). Thus, rLSV closely matched the parental WT LSV from which it was derived and, at this position, matched the published TMA1381 GenBank reference sequence.

To determine whether rLSV retained the biological properties of the WT LSV, we compared replication kinetics and plaque morphology between rLSV and WT LSV. Multistep growth curves in Vero E6 and A549 cells revealed similar replication kinetics, with no appreciable differences in titers (Fig. 1B, C). Consistent with these findings, rLSV produced plaques that were comparable in size to WT LSV in Vero E6 monolayers (Fig. S1A). Together, these data establish a recombinant LSV platform that recapitulates the *in vitro* growth kinetics of the parental virus.

Because UTRs and intergenic regions (IGRs) contain critical regulatory signals for bandavirus transcription and replication, we next mapped the LSV S-segment IGR. The predicted 3′ ends of both the N and NSs mRNAs were located centrally within the IGR and were associated with nearby GC-rich putative transcription termination motifs (Fig. S1B). In the viral (v) RNA orientations shown, the N mRNA 3′ end mapped adjacent to a putative 3′-GGACGG-5′ motif, whereas the NSs mRNA 3′ end mapped adjacent to a distinct putative 3′-GUCCCU -5′ motif. This central IGR-associated arrangement differs from that described for SFTSV, in which the N and NSs mRNA 3′ ends map beyond the IGR and terminate within the coding sequence of the opposing ORF^33^. Thus, the LSV S segment retains the ambisense organization characteristic of bandaviruses but exhibits distinct predicted transcription termination features, providing a genomic framework for future mutagenesis studies using the rLSV platform (Fig. S1C).

### LSV causes rapid, systemic lethal disease in IFNAR^-/-^ mice

Having established that rLSV replicated in mammalian cell lines, including human A549 cells, we next sought to determine whether LSV could cause disease *in vivo*. Type I IFN signaling is a major determinant of susceptibility in related tick-borne bandaviruses, including HRTV and SFTSV^34–36^, therefore, IFNAR^-/-^ mice provide a sensitive model to evaluate viral dissemination and pathogenic potential under conditions of impaired antiviral immunity. To first establish an *in vivo* LSV infection model, we infected IFNAR^-/-^ mice with 10^6^ TCID_50_ of WT LSV. All infected animals rapidly developed clinical signs, including ruffled fur, hunching, lethargy, weight loss, and ocular discharge, and reached humane endpoints^37^ by 4 days post-infection (dpi; Fig. S2A-D). At necropsy, infected animals exhibited gross pathological changes, including pale livers and darkened, congested spleens. Analysis of tissues collected at endpoint revealed widespread viral dissemination, with infectious virus recoverable from liver, spleen, heart, lung, and brain tissues of all WT LSV-infected mice following inoculation of tissue homogenates onto Vero E6 cells (Fig. S2E). Consistent with this broad recovery of infectious virus, high vRNA loads were detected across the same tissues, reaching approximately 10^10^ -10^11^ copies/g in visceral tissues and 10^9^ copies/g in the brain (Fig. S2F). These findings establish that LSV causes rapid, systemic lethal disease in the absence of type I IFN signaling.

Having confirmed the lethality of WT LSV in IFNAR^-/-^ mice, we next evaluated whether rLSV recapitulated this phenotype across a range of inoculation doses. IFNAR^-/-^ mice were infected with 10^5^, 10^3^,10, or 1 TCID_50_ of rLSV and monitored for survival, clinical disease, body weight, and tissue viral burden (Fig. 2). Infection with 10^5^ or 10^3^ TCID_50_ caused rapid disease comparable to WT LSV, with early weight loss, increasing clinical scores, and all animals reaching humane endpoints by 4 dpi (Fig. 2A-L). Infection with 10 TCID_50_ produced a slightly delayed but uniformly lethal disease course (Fig. 2M-P). At the lowest dose tested, 1 TCID_50_, most animals also succumbed to infection, although one mouse survived and exhibited weight gain by 5 dpi (Fig. 2Q-T).

**Figure 2.**
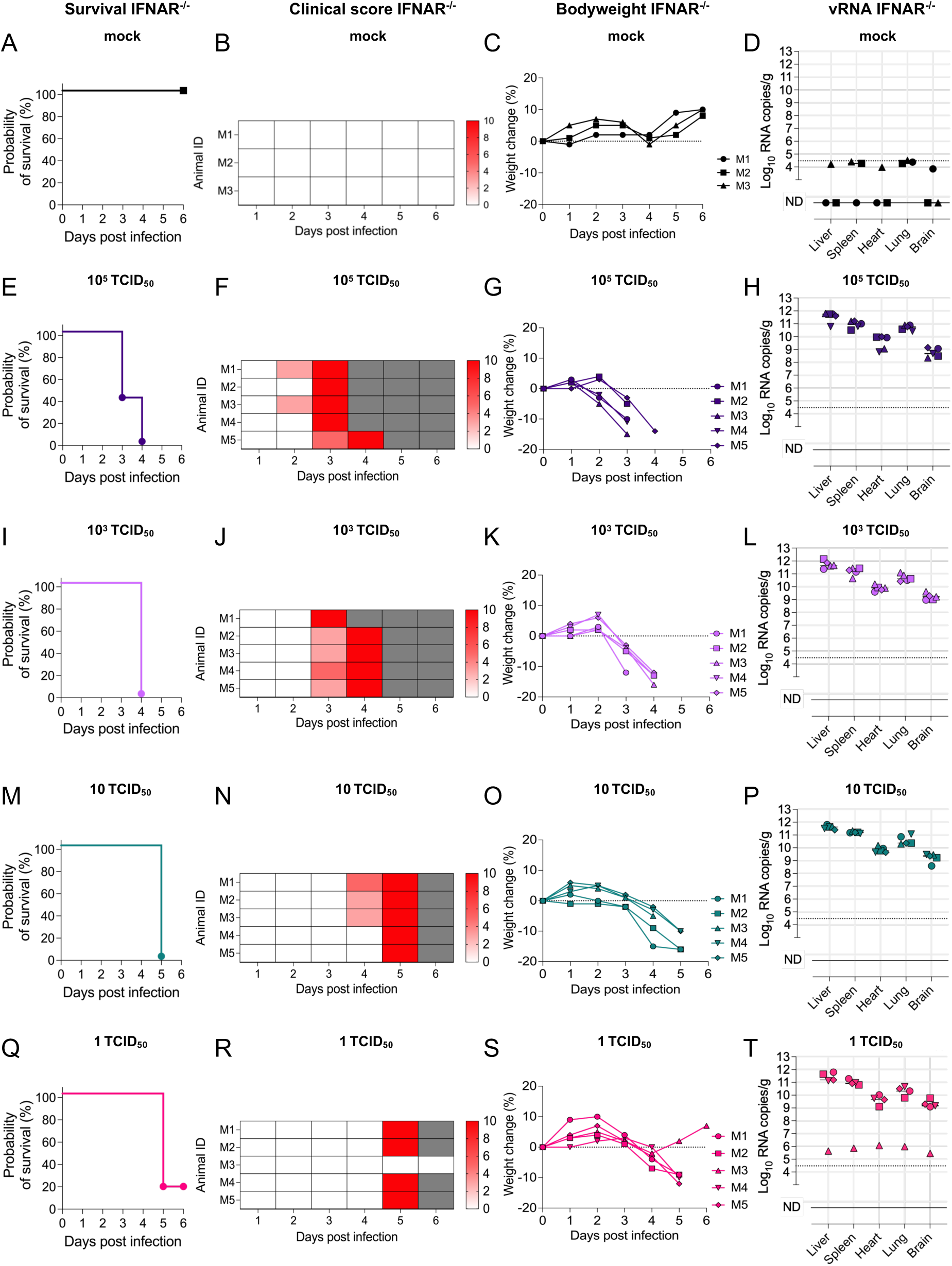
rLSV causes rapid, systemic, and lethal disease in IFNAR^-/-^ mice. IFNAR^-/-^ mice were infected SC with rLSV at doses of 10^5^, 10^3^, 10, or 1 TCID_50_, or mock-infected with Opti-MEM. (A-D) Survival, clinical scores, percent body weight change, and tissue vRNA loads in mock-infected mice. (E-H) Survival, clinical scores, percent body weight change, and tissue vRNA loads in mice infected with 10^5^ TCID_50_ rLSV. (I-L) Survival, clinical scores, percent body weight change, and tissue vRNA loads in mice infected with 10^3^ TCID_50_ rLSV. (M-P) Survival, clinical scores, percent body weight change, and tissue vRNA loads in mice infected with 10 TCID_50_ rLSV. (Q-T) Survival, clinical scores, percent body weight change, and tissue vRNA loads in mice infected with 1 TCID_50_ rLSV. Clinical disease severity is depicted by increasing intensity of red. Animals were euthanized upon reaching humane endpoint. Tissue vRNA loads were measured by RT-qPCR and are shown as log_10_ RNA copies/g tissue. Each symbol represents an individual mouse. Dashed lines indicate the limit of detection. ND, not detected.

Across all lethal rLSV dose groups, high vRNA loads were detected in every animal that reached endpoint and in all tissues examined, including liver, spleen, heart, lung, and brain (Fig. 2H, L, P, T). Because tissues were collected at a humane endpoint rather than at a fixed time point across dose groups, these data reflect terminal viral burden after disease progression. These endpoint tissue burdens were generally in the range of 10^9^ - 10^11^ copies/g across the organs examined, whereas the surviving animal in the 1 TCID_50_ group had substantially lower vRNA levels, approximately 10^5^ - 10^6^ copies/g (M3, diamond symbol; Fig. 2T). Thus, even low-dose rLSV infection resulted in disseminated viral replication and severe disease in IFNAR^⁻/⁻^ mice.

### rLSV infection is associated with hepatic and splenic pathology in IFNAR^-/-^ mice

To assess tissue pathology associated with rLSV infection, hematoxylin and eosin (H&E) staining was performed on liver and spleen sections collected from IFNAR^-/-^ mice infected with either a low (1 TCID_50_) or high (10^5^ TCID_50_) dose of rLSV. Histopathological examination of the liver revealed multifocal hepatocellular necrosis in infected animals, characterized by disruption of normal hepatic architecture, cellular debris, and inflammatory infiltrates (Fig. 3A). These lesions were observed in both low- and high-dose infected mice, whereas liver sections from mock-infected controls displayed preserved lobular architecture with no evidence of necrosis or inflammation. In the spleen, rLSV infection was associated with focal areas of necrosis within the white pulp and disruption of normal lymphoid architecture (Fig. 3B). These changes were evident in infected animals from both dose groups, while spleens from mock-infected controls retained intact lymphoid organization.

**Figure 3.**
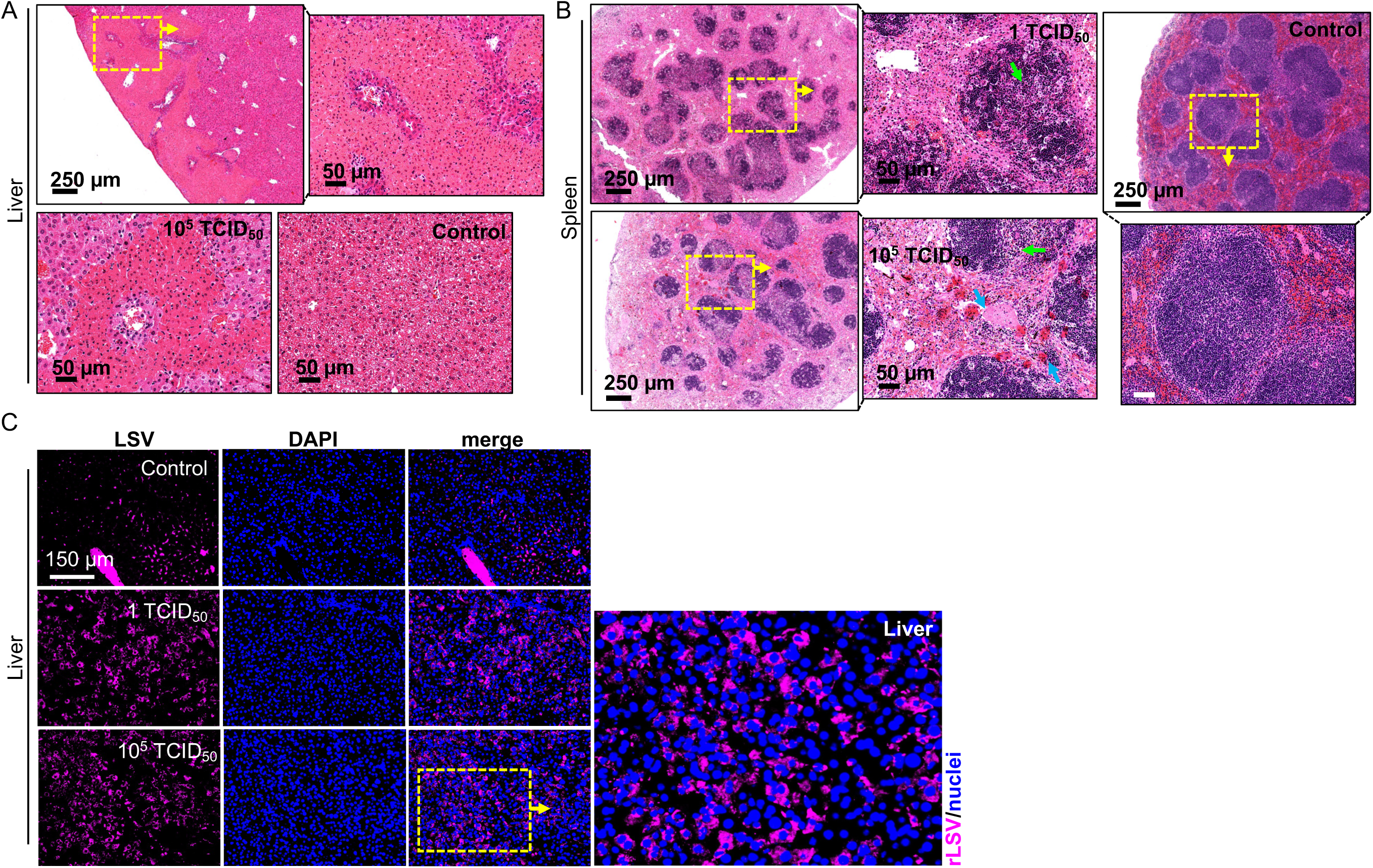
rLSV infection is associated with hepatic and splenic pathology in IFNAR^-/-^ mice. Representative H&E-stained liver and spleen sections from IFNAR^-/-^ mice infected with rLSV at 1 TCID_50_ or 10^5^ TCID_50_ rLSV, or from mock-infected controls. (A) Liver sections showing multifocal hepatocellular necrosis in infected animals, with preserved hepatic architecture in mock-infected controls. Yellow dashed boxes indicate regions shown at higher magnification. (B) Spleen sections showing focal areas of necrosis and disruption of lymphoid architecture in infected animals, compared with intact white pulp organization in mock-infected controls. Yellow dashed boxes indicate regions shown at higher magnification. (C) IF of formalin-fixed liver sections from mock-infected or rLSV-infected IFNAR^-/-^ mice. LSV antigen is shown in magenta, nuclei are stained with DAPI (blue), and merged images are shown. The boxed region is shown at higher magnification in the panel to the right. Scale bars are indicated in each panel.

To determine whether hepatic pathology was associated with viral antigen, immunofluorescence staining was performed on formalin-fixed liver sections using LSV immune ascitic fluid. LSV antigen was detected in liver sections from infected animals, with increased signal in both low-and high-dose infection groups relative to mock-infected controls (Fig. 3C). Together with the high tissue vRNA loads, these findings indicate that systemic rLSV infection in IFNAR^⁻/⁻^ mice is associated with hepatic and splenic pathology, including hepatocellular necrosis and disruption of splenic architecture.

### LSV infection is restricted in immunocompetent C57BL/6J mice

To determine whether LSV causes disease in immunocompetent mice, C57BL/6J mice were infected with 10^5^ TCID_50_ of rLSV and monitored daily for clinical signs and weight loss. Most infected animals remained clinically normal throughout the study, with no evidence of morbidity or mortality (Fig. 4A-E). Consistent with the absence of clinical disease, infected mice maintained or gained weight over time, similar to mock-infected controls (Fig. 4F). To assess viral replication and tissue dissemination, mice were euthanized at 5 or 11 dpi, and vRNA levels were quantified in liver, spleen, heart, lung, and brain by RT-qPCR. No vRNA was detected in any tissues from mock-infected animals (Fig. 4G). In rLSV-infected mice, low levels of vRNA were detected sporadically at both time points, with values near the limit of detection (dashed line) and without consistent tissue tropism (Fig. 4H, I).

**Figure 4.**
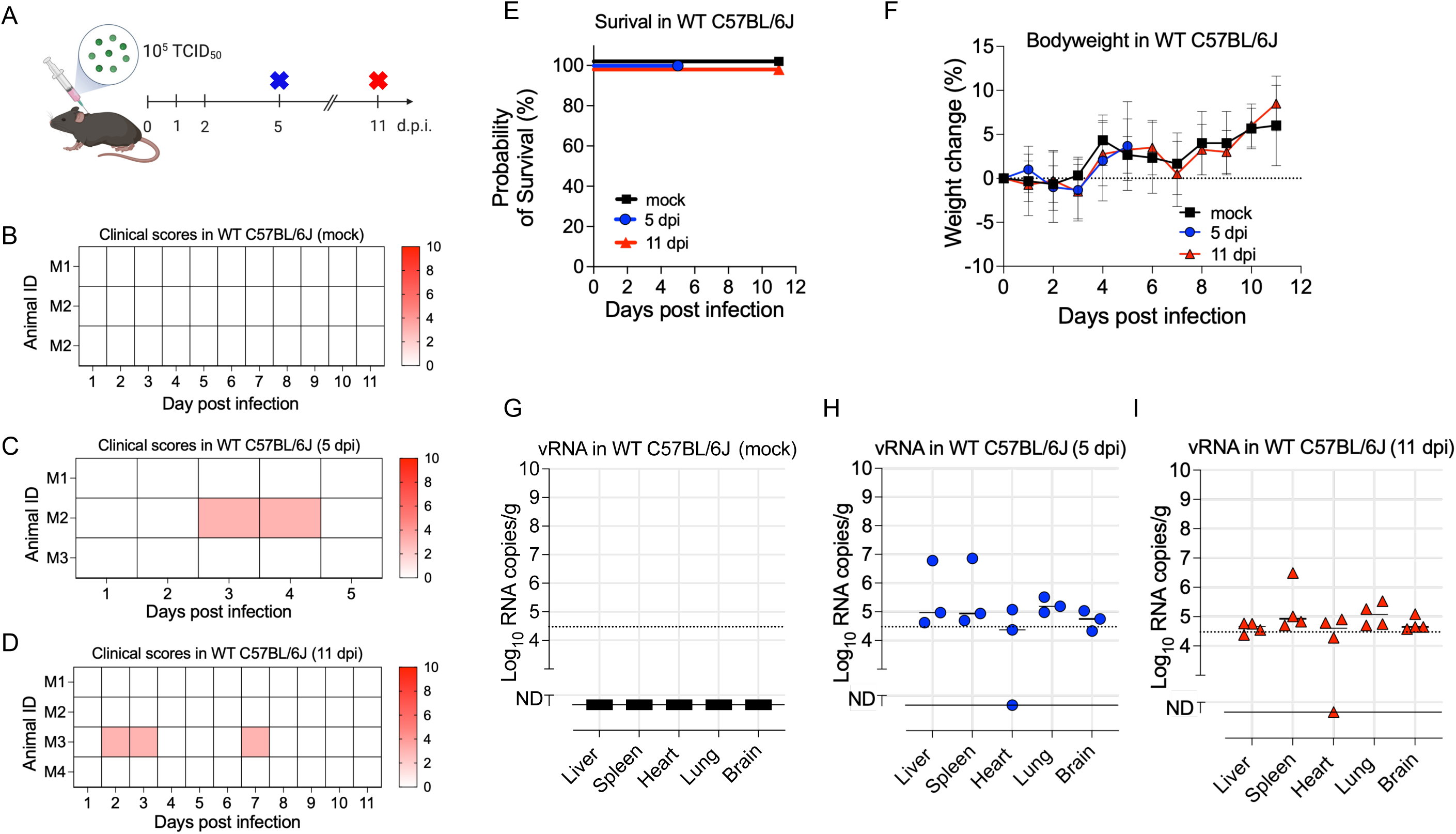
rLSV infection is restricted in immunocompetent C57BL/6J mice. (A) Experimental schematic showing SC inoculation of immunocompetent C57BL/6J mice with 10^5^ TCID_50_ rLSV and serial euthanasia at 5 or 11 dpi. (B-D) Clinical scores of mock-infected mice (B), rLSV-infected mice euthanized at 5 dpi (C), and rLSV-infected mice euthanized at 11 dpi (D). (E) Survival curves showing 100% survival in all groups. (F) Percent body weight change over time in mock-infected and rLSV-infected mice. (G-I) vRNA levels in liver, spleen, heart, lung, and brain tissues from mock-infected mice (G), rLSV-infected mice euthanized at 5 dpi (H), and rLSV-infected mice euthanized at 11 dpi (I), measured by RT-qPCR and shown as log10 RNA copies/g tissue. Dashed lines indicate the limit of detection. ND, not detected. Each symbol represents an individual mouse.

To evaluate systemic inflammatory responses, serum cytokine levels were measured using a multiplex Luminex assay. In contrast to endpoint sera from rLSV-infected IFNAR^-/-^ mice, which exhibited robust induction of IFN-α, IFN-β, IFN-γ, IL-6, and TNF-α, these cytokines were not detected in sera from infected C57BL/6J mice at either 5 or 11 dpi (Fig. 5A-E). Cytokine levels in infected C57BL/6J mice were comparable to those in mock-infected controls. Finally, to determine whether rLSV exposure elicited a humoral immune response, sera collected at 5 and 11 dpi were evaluated for LSV-neutralizing activity. No neutralizing activity was detected at 5 dpi. At 11 dpi, low-level neutralizing activity was detected only at neat or 1:2 serum dilutions (Fig. 5F). These results indicate that immunocompetent C57BL/6J mice restrict rLSV infection, preventing clinical disease, limiting tissue viral burden, and resulting in minimal systemic inflammatory or neutralizing antibody responses.

**Figure 5.**
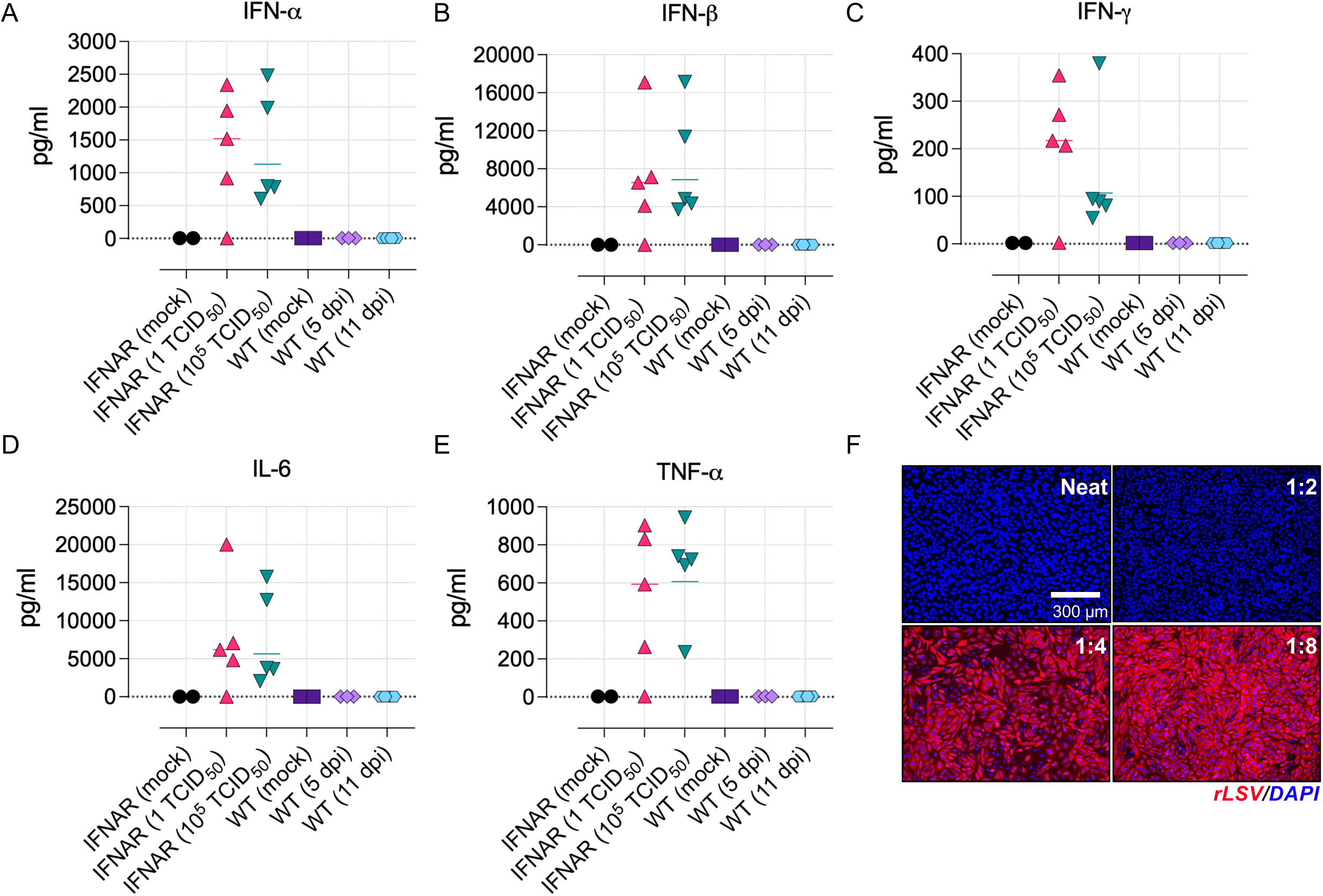
Systemic cytokine responses and neutralizing activity following rLSV infection. (A-E) Serum concentrations of IFN-α (A), IFN-β (B), IFN-γ (C), IL-6 (D), and TNF-α (E) measured by multiplex Luminex assay in mock-infected IFNAR^-/-^ mice, rLSV-infected IFNAR^-/-^ mice infected with 1 or 10^5^ TCID_50_, mock-infected C57BL/6J mice, and rLSV-infected C57BL/6J mice euthanized at 5 or 11 dpi. Cytokine induction was detected primarily in rLSV-infected IFNAR^-/-^mice. Each symbol represents an individual mouse, and horizontal bars indicate the mean. (F) Virus neutralization assay using serum from rLSV-infected C57BL/6J mice collected at 11 dpi. Representative images are shown for neat serum and 1:2, 1:4, and 1:8 serum dilutions. rLSV antigen is shown in red, and nuclei are stained with DAPI (blue). Scale bar, 300 µm.

### Comparative bandavirus infection reveals distinct disease kinetics and limited immune overlap with LSV

To place LSV in the context of clinically relevant tick-borne bandaviruses, we compared rLSV with HRTV strain R124769a and rSFTSV, representing closely related North American and Asian tick-borne bandaviruses, respectively. rSFTSV was generated based on the coding sequences of strain CB8/2016 (genotype D, South Korea; accession numbers: S, MK301488; M, MK301487; L, MK301486). Because complete UTRs were not available for CB8, missing UTR sequences were complemented using the closely related CB1 genotype B strain (accession numbers: S, KY789439; M, KY789436; L, KY789433), enabling successful virus rescue in BSR-T7/5 cells. Phylogenetic analysis of the coding regions confirmed that rLSV, HRTV, and rSFTSV cluster within the genus *Bandavirus* but remain genetically distinct from one another (Fig. 6A). We next compared viral replication in human A549 cells. All three viruses replicated over the course of infection, with rLSV reaching peak titers of approximately 3.5 × 10^5^ TCID_50_/ml, compared with approximately 8.6 × 10^5^ TCID_50_/ml for HRTV and 2.1 × 10^6^ TCID_50_/ml for rSFTSV under these conditions (Fig. 6B).

**Figure 6.**
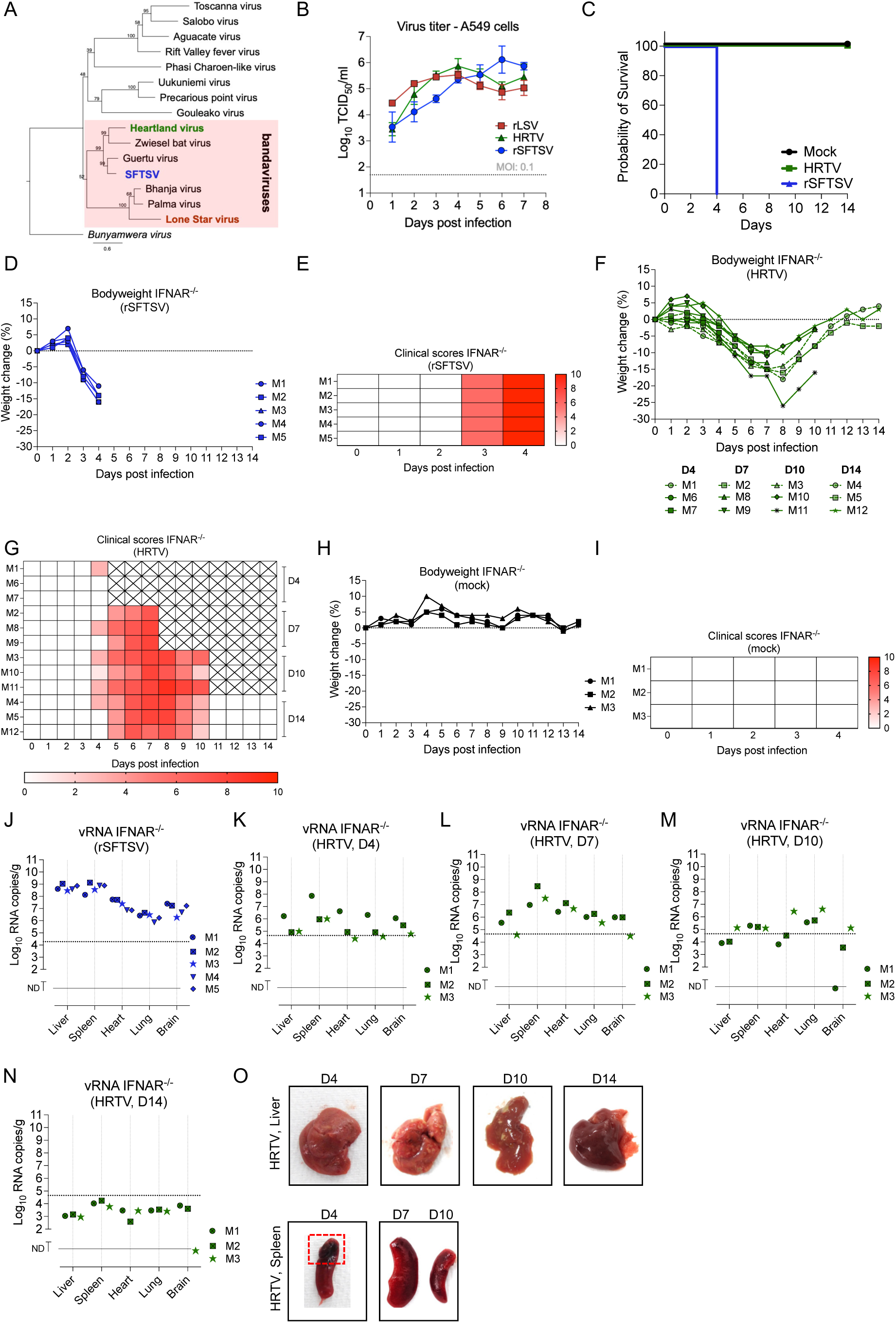
Comparative replication and pathogenesis of tick-borne bandaviruses. (A) Phylogenetic tree showing the relationship of rLSV, HRTV, and rSFTSV within the genus *Bandavirus* relative to other representative bunyaviruses. Bootstrap values are shown at key nodes. (B) Multistep growth kinetics of rLSV, HRTV, and rSFTSV in A549 cells infected at an MOI of 0.1. Supernatants were collected at the indicated time points, and infectious titers were determined by TCID50 assay. Data are shown as mean ± SD (n = 3). (C) Survival curves of IFNAR^-/-^ mice infected with HRTV or rSFTSV, or mock-infected with Opti-MEM. (D) Percent body weight change in rSFTSV-infected mice (E) Clinical scores over time following rSFTSV infection (F) Percent body weight change in HRTV-infected mice (E) Clinical scores over time following HRTV infection (H) Percent body weight change in mock-infected mice. (I) Clinical scores over time following mock infection (J) vRNA levels in tissues of rSFTSV-infected mice at terminal time points. (K-N) vRNA levels in tissues of HRTV-infected mice at 4, 7, 10, and 14 dpi. Tissue vRNA loads were measured by RT-qPCR and are shown as log_10_ RNA copies/g tissue. Dashed lines indicate the limit of detection. ND, not detected. Each symbol represents an individual mouse. (O) Representative gross pathology images of liver and spleen from HRTV-infected mice at the indicated time points. The red dashed box highlights splenic discoloration during acute HRTV infection.

We next compared disease outcomes following infection of IFNAR^-/-^ mice with HRTV or rSFTSV (dose: 10^5^ TCID_50_). Infection with rSFTSV resulted in rapid, uniformly lethal disease, with all animals reaching humane endpoints by 4 dpi (Fig. 6C, D, E). In contrast, HRTV-infected mice developed a sublethal disease course, characterized by transient weight loss and clinical signs, including ruffled fur, hunching, and lethargy, which peaked approximately 6 - 8 dpi before resolving (Fig. 6C, F, G). Mock-infected animals remained clinically normal throughout the experiment (Fig. 6C, H, I). Consistent with these divergent clinical outcomes, rSFTSV-infected mice had high systemic vRNA loads at terminal time points, with the highest burdens detected in liver and spleen, reaching approximately 10^8^–10^9^ copies/g, and lower but readily detectable levels in heart, lung, and brain, generally ranging from approximately 10^6^–10^7^ copies/g (Fig. 6F). HRTV vRNA was also detected across multiple tissues during acute disease, but tissue burdens were more variable between animals and changed over time. The highest HRTV loads were detected in the spleen, reaching approximately 10^6^-10^8^ copies/g at 4 dpi and 10^7^-10^8^ copies/g at 7 dpi, while liver, heart, lung, and brain generally contained lower but detectable vRNA levels during this acute phase. By 10-14 dpi, HRTV vRNA levels declined substantially across most tissues, consistent with clinical recovery and partial viral clearance (Fig. 6K-N). Mock-infected control tissues were negative or near assay background when analyzed using the corresponding virus-specific RT-qPCR assays (Fig. S3). Gross pathological examination of HRTV-infected mice revealed enlarged or discolored livers and spleens during acute disease, with gradual normalization at later time points (Fig. 6O). Together, these findings show that related tick-borne bandaviruses exhibit distinct disease courses in IFNAR^-/-^ mice, with rLSV and rSFTSV causing rapid lethal disease, whereas HRTV establishes a sublethal infection under the conditions tested.

Finally, we asked whether immunity induced by HRTV infection protected against homologous or heterologous bandavirus challenge. IFNAR^-/-^ mice were first infected with HRTV and monitored through clinical recovery, then rechallenged with HRTV, rSFTSV, or rLSV (Fig. 7A). Primary HRTV infection caused transient weight loss and moderate clinical disease, followed by recovery from all HRTV-associated disease (Fig. 7B-H). One HRTV-infected mouse was euthanized before rechallenge because of a non-infection-related injury (M5, Fig. 7B, C). Upon homologous rechallenge with HRTV at 28 dpi, mice showed no weight loss or clinical signs, consistent with protective immunity against HRTV (Fig. 7B, C). HRTV-immune mice challenged with rSFTSV were also protected from severe disease, exhibiting minimal weight loss and no clinical signs despite the lethality of rSFTSV in naïve IFNAR^-/-^ mice (Fig. 7E, F). In contrast, HRTV-immune mice challenged with rLSV developed rapid weight loss and severe clinical disease, requiring euthanasia (Fig. 7G, H). These results indicate that immunity induced by primary HRTV infection protects IFNAR^-/-^ mice against homologous HRTV rechallenge and heterologous rSFTSV challenge but is insufficient to protect against rLSV. Together, these findings support functional differences in protective immunity among related tick-borne bandaviruses and indicate that HRTV-induced immune responses do not confer comparable protection against LSV under these conditions.

**Figure 7.**
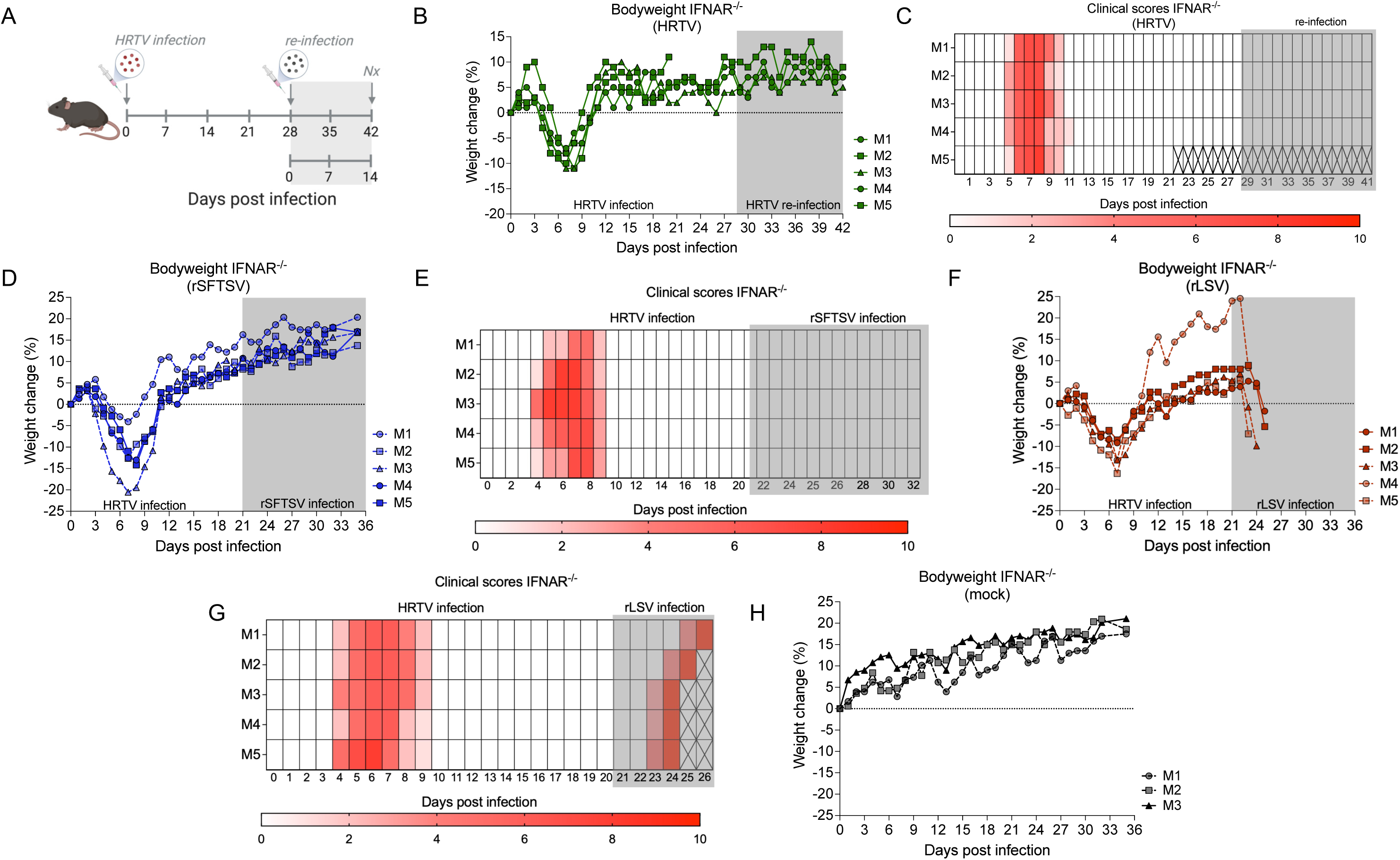
HRTV-induced immunity protects against HRTV and rSFTSV but not rLSV in IFNAR^-/-^ mice. (A) Experimental schematic illustrating primary HRTV infection followed by homologous HRTV rechallenge or heterologous challenge with rSFTSV or rLSV. (B, C) Percent body weight change (B) and clinical scores (C) during primary HRTV infection and homologous HRTV rechallenge. (D, E) Percent body weight change (D) and clinical scores (E) during primary HRTV infection and heterologous rSFTSV rechallenge. (F, G) Percent body weight change (F) and clinical scores (G) during primary HRTV infection and heterologous rLSV rechallenge. (H) Percent body weight change in mock-infected control mice. Clinical disease severity is depicted by increasing intensity of red. Shaded regions indicate the rechallenge or heterologous challenge period. Cross-hatched boxes indicate time points after animals were euthanized or were no longer monitored. Each line represents an individual mouse.

## Discussion

LSV has remained largely uncharacterized despite its close phylogenetic relationship to pathogenic tick-borne bandaviruses^5,14,15,20,38,39^. The recent detection of LSV RNA in cerebrospinal fluid from an immunocompromised patient with fatal meningoencephalitis^24^ highlights the need for experimental systems to evaluate LSV’s pathogenic potential. Here, we establish a reverse genetics system for LSV, define its infection phenotype in immunocompromised and immunocompetent mouse models, and compare LSV with the related tick-borne bandaviruses HRTV and SFTSV.

Our findings demonstrate that LSV replication is heavily restricted by type I IFN signaling in mice. In IFNAR^-/-^ mice, both WT LSV and rLSV caused rapid, systemic, and lethal disease, with high viral burdens across multiple tissues and pathological changes in the liver and spleen. In contrast, infection was efficiently controlled in immunocompetent C57BL/6J mice, which showed no clinical disease, only sporadic low-level vRNA detection, no recovery of infectious virus, minimal systemic cytokine induction, and weak neutralizing activity. This pattern is consistent with prior studies showing that type I IFN signaling is a major determinant of susceptibility to pathogenic bandaviruses, including SFTSV^34^ and HRTV^35,36^. Thus, rather than identifying type I IFN restriction as unique to LSV, our data extend this established principle to a previously neglected member of the genus and provide the first *in vivo* framework for studying LSV pathogenesis.

The development of a reverse genetics system for LSV is a central advance of this study. Recombinant virus systems have been essential for defining bandavirus replication, virulence, and immune evasion mechanisms, including studies of HRTV and SFTSV NSs-mediated antagonism of innate immune signaling^7,8,28,32,33^. For HRTV, reverse genetics has enabled direct testing of NSs function and demonstrated that NSs contributes to virulence and attenuation^35^. Until now, comparable tools were not available for LSV, limiting mechanistic investigation of this virus. We show that rLSV recapitulates the *in vitro* replication properties and *in vivo* disease phenotype of the parental virus, establishing a genetically tractable platform for future studies of LSV replication, reporter-virus development, and viral determinants of pathogenesis. Because LSV can be studied under BSL-2 conditions, this system may also provide a complementary lower-containment model for examining conserved and divergent features of tick-borne bandavirus biology, while recognizing that LSV is biologically distinct and should not be viewed as a direct substitute for HRTV or SFTSV.

Comparative analysis with HRTV and rSFTSV further showed that related bandaviruses differ in disease kinetics and outcomes *in vivo*, despite shared classification and overlapping replication competence in cell culture. In A549 cells, rLSV, HRTV, and rSFTSV all replicated over the course of infection, although rLSV reached somewhat lower peak titers under the conditions tested. In IFNAR^-/-^ mice, however, rLSV and rSFTSV caused rapid, lethal disease, whereas HRTV produced a sublethal infection characterized by overt clinical signs, weight loss, and vRNA detection, followed by recovery. Our HRTV phenotype is consistent with prior HRTV animal models, demonstrating that HRTV lethality is dependent on inoculation route and the extent of IFN pathway disruption^36^. Nevertheless, under the conditions used here, the three viruses occupied distinct positions along a spectrum of disease severity. This reinforces the idea that phylogenetic relatedness alone does not predict pathogenic outcome. Instead, differences in viral antagonism, replication kinetics, tissue tropism, and dissemination likely contribute to disease outcome, although the specific viral determinants responsible for these phenotypes remain to be defined.

The rechallenge studies also revealed functional differences in protective immunity among these viruses. Prior HRTV infection protected IFNAR^-/-^ mice against homologous HRTV rechallenge and against otherwise lethal rSFTSV challenge, consistent with previous studies showing cross-protective immunity between HRTV and SFTSV^30,40,41^. In contrast, HRTV-induced immunity was insufficient to protect against rLSV challenge under the same conditions. This result does not define the precise antigenic relationship among these viruses, and the low neutralizing titers generated after rLSV exposure in C57BL/6J mice limit conclusions about LSV antigenicity. However, the failure of HRTV-induced immunity to protect against rLSV supports functional differences in protective immune responses and suggests that immune determinants shared between HRTV and SFTSV are absent, poorly conserved, or insufficiently protective in the context of LSV challenge. Future studies using higher-titer immune sera, monoclonal antibodies, and defined glycoprotein immunogens will be needed to map the antigenic and protective relationships among LSV, HRTV, and SFTSV more precisely.

These findings have broader relevance in the context of changing tick ecology and emerging bandavirus risk in North America. *A*. *americanum*, the Lone star tick, is an aggressive human-biting tick with an expanding geographic range and is implicated in transmission or maintenance of multiple pathogens, including HRTV^42–46^. The presence of poorly characterized bandaviruses in expanding tick populations emphasizes the importance of studying endemic viruses before their clinical significance is fully apparent. At the same time, the establishment of additional tick species, including *Haemaphysalis longicornis*, in the U.S. raises concern about future changes in tick-borne virus ecology^47,48^. While the likelihood of emergence, spillover, or reassortment^49^ among bandaviruses remains uncertain and will depend on vector competence, host range, geographic overlap, and co-infection dynamics. Defining the replication, pathogenic, and immunological properties of endemic viruses such as LSV is important for surveillance and preparedness.

In summary, this study establishes LSV as a genetically tractable, BSL-2-accessible bandavirus that is strongly restricted by type I IFN signaling but capable of causing rapid systemic disease in immunocompromised mice. By integrating reverse genetics, mouse models, and comparative infection studies with HRTV and rSFTSV, we provide a foundation for future mechanistic studies of LSV biology and for broader investigation of conserved and divergent determinants of bandavirus pathogenesis.

## Materials and Methods

### Ethics statement

All animal studies were conducted in accordance with protocols approved by the Indiana University School of Medicine (IUSM) Institutional Animal Care and Use Committee (IACUC; protocol #22080). Experiments involving LSV were performed in BSL-2 or Animal BSL-2 (ABSL-2) facilities, while experiments with SFTSV and HRTV were conducted in IUSM’s BSL-3 and ABSL-3 facilities.

### Cells and viruses

Vero E6 cells (African green monkey kidney cells), BSR-T7/5 cells (baby hamster kidney cells), which stably express T7 RNA polymerase^50^ maintained with 1 mg/ml of G418 (Geneticin; Invitrogen) and A549 (human alveolar adenocarcinoma epithelial cells) were grown in Dulbecco’s modified Eagle medium (DMEM; Gibco) supplemented with 2 – 10% (V/V) fetal bovine serum (Gibco). All cells were grown at 37°C and 5% CO_2_.

Lone star virus TMA 1381 (i.e., WT LSV) was obtained from the World Reference Collection of Emerging Viruses and Arboviruses (WRCEVA) at the University of Texas Medical Branch (UTMB), passaged in Vero E6 cells once, and plaque-purified to create stocks. Stocks were then sequenced via Sanger sequencing (ACGT) using segment-specific primers (Supplemental Table 1). HRTV isolate R124769a was obtained from the CDC Arbovirus Reference Collection (originated from a human serum sample collected in Kentucky in 2017). All virus stocks were grown and titrated in Vero E6 cells by 50% tissue culture infectious dose assay (TCID_50_).

### Recombinant LSV and SFTSV generation

Full-length rescue plasmids for rLSV were generated by amplifying positive-sense cDNA corresponding to the S, M, and L genome segments from LSV TMA 1381 (WT LSV). These cDNA amplicons were assembled using Gibson Assembly (NEBuilder HiFi DNA Assembly, NEB) into plasmids containing a T7 promoter upstream and a hepatitis delta virus ribozyme followed by a T7 terminator downstream (pUC plasmid, IDT), yielding the plasmids pT7LSV-S, pT7LSV-M, and pT7LSV-L. To generate rSFTSV, the positive-sense coding regions of the S, M, and L segments from strain CB8/2016 (GenBank accession numbers: S, MK301488; M, MK301487; L, MK301486) were combined with UTR sequences from strain CB1 (GenBank accession numbers: S, KY789439; M, KY789436; L, KY789433). These sequences were synthesized as gBlocks Gene Fragments (IDT) and assembled into T7-driven rescue plasmids using Gibson Assembly as described above. Helper plasmids encoding the N and RdRp for each virus were generated by cloning the corresponding coding regions into pTM1 vectors^51^ under the control of a T7 promoter and encephalomyocarditis virus (EMCV) internal ribosome entry site (IRES), as previously described^52^. These constructs were designated pTM1-N and pTM1-L. Recombinant viruses were rescued by transfecting BSR-T7/5 cells with 1 µg each of the full-length S, M, and L segment plasmids together with 0.5 µg of each helper plasmid using TransIT-VirusGEN Transfection Reagent (Mirus Bio) in 6-well plates. Seven days post-transfection, rescues were passaged onto Vero E6 cells. Successful virus rescue was confirmed by the appearance of cytopathic effect (CPE) observed under phase-contrast microscopy and by immunofluorescence (IF) using mouse immune ascitic fluids (MIAF; WRCEVA, UTMB) for LSV and an anti-N SFTSV antibody (kind gift from Prof. Paul Duprex, University of Pittsburgh) for SFTSV. Virus stocks were subsequently amplified and titrated in Vero E6 cells.

### Viral growth kinetics

Vero E6 and A549 cells (2.0 x 10^5^ cells/ml, 24-well plate) were infected with either LSV, rLSV, HRTV or rSFTSV at a multiplicity of infection (MOI) 0.1 for 1 h at 37°C. Cell monolayers were washed 3X with D-PBS (Gibco) and then provided with DMEM/2% FBS. At the desired time points, supernatant was collected, and infectious viral titers were determined by TCID_50_ assay.

### TCID_50_ and plaque assays

TCID_50_ assays were performed in Vero E6 cells at a density 10^4^ cells per well in 96-well plates and infected with a 10-fold serial dilution of virus. At 7 d.p.i., CPE was recorded and viral titers were expressed as TCID_50_ units/ml. Values were calculated using the Reed-Muench method. Plaque assays were performed in 6-well plates with Vero E6 cells at a density of 10^5^ cells/ml. Cell monolayers were infected with a 200 μl inoculum of virus diluted in Opti-MEM. Cells were then overlaid with 0.6% Avicel (FMC, Avicel RC-591) in 2X minimum essential medium (MEM)/2% FBS. Cells were fixed at 7 days post-infection (d.p.i) with 4% (w/v) paraformaldehyde (PFA) in PBS for 30 minutes, and plaques were visualized using crystal violet.

### Quantitative RT-qPCR

LSV, rLSV, HRTV, and rSFTSV vRNA were quantified using primers targeting the S genome segment (Supplemental Table 1). Primers and probes were used at working concentrations of 10 μM and 5 μM, respectively. To generate RNA for a standard curve, pT7-S plasmids containing the entire S segments were linearized via restriction enzyme digestion, purified, and used as a template for *in vitro* RNA transcription (MEGAscript T7 transcription kit; Invitrogen). The resulting RNA was then diluted to known copies per ml in RNase-free water and serially diluted for each assay. RT-qPCR was performed using the Luna Universal Probe One-Step RT-qPCR Kit (NEB) on the QuantStudio^™^ 5 (ThermoFisher Scientific) using the following conditions: 55°C for 10 minutes for reverse transcription, followed by 95°C for 1 minute, and then 40 cycles of 95°C for 10 seconds and 60°C for 1 min. For the animal studies, RNA levels were reported as log vRNA copies per gram of tissue, with the limit of detection (LoD) calculated based on the highest cycle threshold (Ct) value on the standard curve for each assay.

### Animal study design

Six- to eight-week-old female IFNAR^-/-^ mice (The Jackson Laboratory) and immunocompetent C57BL/6J mice were housed in HEPA-filtered cages in either ABSL-2 or ABSL-3 facilities with *ad libitum* access to food and water. All animal experiments were conducted in accordance with approved IACUC protocols. In the initial experiment assessing LSV pathogenicity, (n = 5) IFNAR^-/-^ mice were subcutaneously (SC) infected with 100 µL of plaque-purified WT LSV at a dose of 10^6^ TCID_50_ diluted in Opti-MEM, while (n = 3) control mice received 100 µL of Opti-MEM alone. Mice were monitored daily for clinical signs of disease, and body weights were recorded. A dose-response study was subsequently performed in IFNAR^-/-^ mice. Groups of five mice (n = 5) per dose were SC infected with 100 µL of rLSV at doses of 10^5^, 10^3^, 10, or 1 TCID_50_ diluted in Opti-MEM. A control group (n = 3) of mice received 100 µL of Opti-MEM alone. Mice were monitored daily for clinical signs and weight loss and were euthanized upon reaching predetermined humane endpoints based on a standardized clinical scoring system, as we have previously described^37^. To assess LSV infection in immunocompetent hosts, C57BL/6J mice were SC infected with 100 µL containing 10^5^ TCID_50_ rLSV diluted in Opti-MEM and serially euthanized at 5 dpi (n = 3) or 11 dpi (n = 4) with infected mice per time point. Control animals (n = 3) received 100 µL of Opti-MEM alone. Mice were monitored daily for clinical signs and weight loss. At the time of euthanasia, mice were anesthetized with isoflurane and terminally exsanguinated for blood collection. Following cervical dislocation, tissues including liver, spleen, heart, lungs, and brain were harvested for viral isolation and RNA extraction, followed by RT-qPCR analysis of vRNA. To assess the pathogenesis of HRTV and rSFTSV in IFNAR^-/-^ mice, animals were SC infected with 100 µL containing 10^5^ TCID_50_ of either HRTV or rSFTSV and monitored as described above. For rSFTSV infection, mice (n = 5) were infected and euthanized upon reaching the humane endpoint. For HRTV infection, groups of three mice per time point (i.e., n = 3) were euthanized at 4, 7, 10, and 14 dpi for longitudinal assessment of viral burden and gross pathology. Mock-infected control mice (n = 3) received 100 µL of Opti-MEM alone. To assess immune cross-protection between bandaviruses, an *in vivo* rechallenge experiment was performed in IFNAR^-/-^ mice. Three groups of 5 mice each were first SC infected with 100 µL containing 10^5^ TCID50 HRTV and monitored daily for clinical signs and weight loss. Mice were then rechallenged at 28 dpi with 100 µL containing 10^5^ TCID50 of HRTV, rSFTSV, or rLSV (3 groups, n = 5). Mice were monitored daily for clinical signs and weight loss following rechallenge.

### Virus Neutralization Assay

The serum obtained from rLSV-infected C57BL/6J mice was aliquoted and heat-inactivated at 56°C for 30 minutes in a water bath. Two-fold dilutions of the inactivated serum were mixed with 10^2^ TCID_50_ rLSV and incubated at 37°C for 1 hr. Fifty µL of the serum and virus mixture was then used to infect Vero E6 cells in a 96-well plate and incubated at 37°C for 72 hrs. The plates were then fixed in 4% paraformaldehyde (PFA) and stained with 50 µL of 1:1000 LSV MIAF primary antibody and incubated at 4°C overnight on a shaker, washed, and stained with 50 µL of 1:1000 diluted Alexa Fluor 594 goat anti-mouse IgG (H&L) secondary antibody (Invitrogen) for 1 hour in the dark at room temperature. The plates were then washed, and 50 µL of 1:1000 of 4′, 6-diamidino-2-phenylindole (DAPI) nuclear stain (Fisher EN2248) for nucleus staining was added and incubated at room temperature for 5 minutes before washing. The plates were then observed using a fluorescence microscope (EVOS M5000 imaging system, ThermoFisher). Neutralization was assessed by reduction or absence of LSV antigen-positive cells relative to virus-only control wells.

### Histological examination

Liver, spleen, and brain tissue samples from rLSV-infected IFNAR^-/-^ and Opti-MEM-treated IFNAR^-/-^ mice were collected and fixed in 4% PFA for 24 hrs, followed by 100% EtOH, then embedded in paraffin. The formalin-fixed paraffin-embedded (FFPE) tissue sections were stained with hematoxylin and eosin (H&E) for histological examination. All the processing, including embedding, sectioning, and staining, was done by the IUSM histology services core.

### Immunofluorescence (IF)

rLSV-infected and uninfected FFPE liver sections were de-paraffinized and hydrated in HistoClear and decreasing concentrations of ethanol (100 – 50%). The heat-induced antigen retrieval method was used to retrieve the antigen, and the slides were then stained with 30 µL of a 1:300 LSV MIAF primary antibody and incubated at 4°C overnight, washed, and stained with 30 µL of a 1:200 Alexa 594 anti-mouse secondary antibody for 1 hour in the dark at room temperature. The slides were then washed, ProLong Gold mounting media added, and then observed using a fluorescence microscope (EVOS M5000 imaging system, ThermoFisher).

### Cytokine analysis

Whole blood samples from mice were collected via cardiac puncture and placed in SST microtainer tubes (BD). Serum was separated by centrifugation at 2,000 g for 10 min. IFN-α, IFN-β, IFN-γ, IL-6, and TNF-α were detected and quantified using the ProcartaPlex™ Mouse Simplex and Combinable Panels (Invitrogen). The assay was performed according to the manufacturer’s recommendation, and cytokine expression was measured on a Luminex-200 system and the XMap Platform (Luminex Corporation, Austin, TX, USA).

## Statistical Analysis

Data were analyzed and plotted using GraphPad Prism 10. Survival curves were compared using the log-rank Mantel-Cox test. Viral titers, vRNA loads, cytokine concentrations, clinical scores, and body weight changes are presented descriptively as individual animals and/or mean values with error bars as indicated in the figure legends. A *P* value of <0.05 was considered statistically significant.

## Data availability

All relevant data are included in the manuscript and its supporting information files.

## Acknowledgments

We thank the Laboratory Animal Resource Center (LARC) staff at Indiana University School of Medicine (ISUM) for assisting with mouse husbandry and care throughout this study. We also thank Dr. George Sandusky (Department of Pathology, IUSM) for histology consultation. This work was funded by a CTSI Biomedical Research Grant and IUSM start-up funds.

---

**Supplemental Figure 1. Plaque morphology and S-segment transcription termination mapping for LSV.** (A) Representative plaque morphology of WT LSV and rLSV on Vero E6 cells at the indicated dilutions. (B) Mapping of putative transcription termination motifs within the LSV S-segment IGR. The S-segment antigenome and genome orientations are shown with the N and NSs ORFs indicated. Representative Sanger sequencing traces of N and NSs mRNAs are shown beneath each map. Dotted vertical lines indicate the predicted 3′ ends of the N and NSs mRNAs, and highlighted sequences indicate nearby putative transcription termination motifs within the IGR. (C) Schematic representation of the LSV S-segment ambisense coding strategy and the relative positions of the N and NSs mRNAs, illustrating that both mapped mRNA 3′ ends are located within the IGR.

**Supplemental Figure 2. WT LSV causes rapid systemic disease in IFNAR^-/-^ mice.** IFNAR^-/-^mice were infected SC with 10^6^ TCID_50_ WT LSV or mock-infected with Opti-MEM. (A) Survival curve of WT LSV-infected and mock-infected mice. (B) Clinical scores of mock-infected and WT LSV-infected mice. Clinical disease severity is depicted by increasing intensity of red. (C, D) Percent body weight change in mock-infected (C) and WT LSV-infected (D) mice. (E) Number of mice from which infectious virus was recovered from liver, spleen, heart, lung, and brain at necropsy. (F) vRNA loads in liver, spleen, heart, lung, and brain tissues from WT LSV-infected and mock-infected mice, measured by RT-qPCR and shown as log_10_ RNA copies/g tissue. The dashed line indicates the limit of detection. ND, not detected. Each symbol represents an individual mouse.

**Supplemental Figure 3. Mock-infected control tissues analyzed using virus-specific RT-qPCR assays.** (A) vRNA levels in mock-infected IFNAR^-/-^ mouse tissues were analyzed using the SFTSV-specific RT-qPCR assay. (B) vRNA levels in mock-infected IFNAR^-/-^ mouse tissues were analyzed using the HRTV-specific RT-qPCR assay. Tissue vRNA loads were measured by RT-qPCR and are shown as log_10_ RNA copies/g tissue. Dashed lines indicate the limit of detection. ND, not detected. Each symbol represents an individual mouse.

**Supplementary Table 1. Oligonucleotides used in this study.** Primer and probe sequences used for LSV genome sequencing, S-segment RACE, and virus-specific RT-qPCR assays for LSV, HRTV, and SFTSV. Primer/probe names, target virus, genome segment, sequence, and intended purpose are indicated.

